# Regulation of polarity during morphological transitions in *Aureobasidium pullulans*

**DOI:** 10.64898/2026.10.16.759656

**Authors:** Clara Fikry, Avril Wang, Adryanne Gonzalez, Daniel J. Lew

## Abstract

Many fungi are able to switch between different morphologies by regulating the behavior of the cytoskeleton. In turn, the cytoskeleton is regulated by Rho-family GTPases, whose differing spatiotemporal patterns of activity lead to different modes of growth. The polymorphic fungus *Aureobasidium pullulans* alternates between multiple growth modes, including multibudding, isotropic, and hyphal growth. Here, we describe hyphal growth and the yeast-to-hyphal transition in this organism, with a focus on cell polarity. Yeast cells switched between cycles of budding and isotropic expansion in a variable manner prior to initiating hyphal extension. Initial hyphal extension proceeded in a stop-and-start manner with frequent pauses, before switching to continuous growth. Hyphal branching occurred from the sides of hyphae, and septation at multiple sites occurred in more mature hyphae. Previous work showed that either of two Rho-family GTPases, Rac1 and Cdc42, were sufficient promote budding. However, while Rac1 was also able to drive hyphal extension without Cdc42, Cdc42 could not promote hyphal growth without Rac1. Our findings provide the basis for an investigation of how polarity components are re-purposed for different fungal growth modalities.

## INTRODUCTION

Cells come in an amazing variety of shapes. Cell shape is determined by the dynamic organization of the cytoskeleton. In animals, the forces provided by cytoskeletal structures directly deform the plasma membrane to change cell shape (Fletcher & Mullins, 2010). However, plant and fungal cells are encased in a cell wall and inflated by turgor pressure resulting from water influx (Harold, 2002; Money, 1990). In these cells, the cytoskeleton controls cell shape by governing where and when the cell wall is remodeled, allowing turgor to drive directed expansion and polar growth (Lew, 2011).

Fungal cells proliferate predominantly in one of two morphologies: yeasts or hyphae. Cell wall modification is mediated by local exocytosis of vesicles carrying cell wall synthases and hydrolases, which are delivered to designated sites by cytoskeletal tracks (Riquelme et al., 2018). Growth zones are specified by a network of polarity factors centered on a conserved subfamily of Rho GTPases called Cdc42 and Rac (Arkowitz & Bassilana, 2015; Etienne-Manneville, 2004; H.-O. Park & Bi, 2007). These GTPases cycle between active GTP-bound states and inactive GDP-bound states. GTP-bound Cdc42 and Rac interact with various effectors to orient the cytoskeleton and promote the delivery of secretory vesicles to polarity sites (Arkowitz & Bassilana, 2015; Chiou et al., 2017). Thus, fungal cell morphology depends on where and when these GTPases become active.

In yeasts, different spatiotemporal patterns of Cdc42 activity can lead to budding (as in *Saccharomyces cerevisiae*) or polar extension (as in *Schizosaccharomyces pombe*). *S. cerevisiae* cells are ovoid and polarize Cdc42 to a single patch of the cortex, leading to bud emergence at that site (Bi & Park, 2012). Subsequent depolarization of Cdc42 within the bud allows for isotropic bud expansion. *S. pombe* cells are cylindrical and polarize Cdc42 to the two ends of the cell, directing bipolar extension (Chiou et al., 2017). Recent work has shown that yeast cells of the multibudding *Aureobasidium pullulans* become multinucleate and polarize Cdc42 and Rac to a variable number of cortical sites, leading to the simultaneous formation of multiple daughter cells (Crocker et al., 2025). Thus, a major determinant of yeast cell morphology is the number and location of polarity sites that can form.

Filamentous fungi grow hyphae (long thin tubes) by focusing Cdc42 and/or Rac to a single tip for a prolonged time (Arkowitz & Bassilana, 2015). Hyphal extension can continue for many cell cycles without division, leading to multinucleate hyphae, but hyphae can also be partitioned by septa (Riquelme et al., 2018). Hyphae also grow branches by forming additional polarity sites at the sides or tips of existing hyphae, and in many species the branched hyphal networks can become more interconnected by fusion of hyphae (Harris, 2019; Read et al., 2014). More elaborate structures, such as those used to trap nematode worms by carnivorous fungi, or those used to make fruiting bodies like mushrooms for spore dispersal, are built by turning and interleaving hyphae in complex programmed patterns (Kriegler et al., 2025; Riquelme et al., 2018). Thus, a major determinant of hyphal cell morphology is the temporal change in the direction of growth at individual polarity sites.

Dimorphic fungi are capable of shifting between yeast and hyphal forms (Boyce & Andrianopoulos, 2015; Nigg & Bernier, 2017). Hyphae generally emerge from germinating spores or from yeast cells. In turn, yeast cells can bud off from hyphae (Oujezdsky et al., 1973; Ramos & García Acha, 1975; Salazar & Restrepo, 1985). For many species, switching between growth modes is linked to the infection cycle (cues from their host) or to the sexual cycle (pheromones) (Boyce & Andrianopoulos, 2015; Nigg & Bernier, 2017; Riquelme et al., 2018). Hyphal-inducing signals include changes in temperature, nutrient status, pH, and serum. Such signals initiate signal transduction cascades and transcriptional changes that promote hyphal development (Chow et al., 2021; Kinnaer et al., 2019; Nigg & Bernier, 2017; Riquelme et al., 2018; Ruiz-Herrera et al., 1995; Zhao et al., 2019).

*A. pullulans* is a polymorphic fungus that can grow by single budding or multi-budding, by branching hyphal growth, by isotropic expansion without budding, or by meristematic division (Goshima, 2022; Mitchison-Field et al., 2019; Ramos & García Acha, 1975; Slepecky & Starmer, 2009; Wirshing & Lew, 2026). Yeast-to-hypha transitions occur readily in lab conditions, but the signals that induce the switch and the cell biological events that accompany the switch have not been elucidated. In this work, we describe the process of hyphal initiation from yeast cells in *A. pullulans*, with a focus on the behavior of polarity factors. Hyphal outgrowth was preceded by a variable number of budding or isotropic expansion cycles. Hyphal initiation was probabilistic, with cells being more likely to initiate a hypha as they increased in nuclear number. Unexpectedly, early hyphal growth was intermittent, characterized by frequent pausing involving temporary depolarization. Thus, rather than a deterministic switch, the cells of *A. pullulans* can select one of several options for how to deploy their polarity circuit, in a somewhat stochastic manner. These findings set the stage for understanding how the same polarity circuit is regulated so as to yield different polarity outcomes in successive cell cycles.

## RESULTS

### Localization of polarity factors during hyphal growth

Various factors including nutrients and cell density have been shown to affect the mode of growth in *A. pullulans* (Finlay, 1987; Goshima, 2022; D. Park, 1984; Ramos & García Acha, 1975; Vinroot & Torzilli, 1988). We used low cell densities to induce hyphal growth in microscopy experiments (see methods). As hyphae matured, we observed septation, branching, and budding from hyphae (Supplemental Movie 1). Septa typically formed at locations distant from growing tips. We did not observe hyphal fusion under our experimental conditions.

Previous work identified Rac1 and Cdc42 as core polarity GTPases in this fungus, and the p21-activated kinase Pak1 as a Rac1 effector that equalizes polarity sites in multibudding cells (Crocker et al., 2025). To assess the localization of these polarity factors in hyphae, we tagged endogenous polarity proteins with fluorescent proteins codon-optimized for *A. pullulans*. Strains bearing the tagged proteins did not display growth defects (Supplemental Figure S1A). However, tagging of endogenous Rac1 with mNeonGreen (mNG) led to reduced abundance of hyphae at colony edges (Supplemental Figure S1B-D) and a slight widening of hyphae, indicating that this probe is not fully functional (Figure 1).

**Figure 1.**
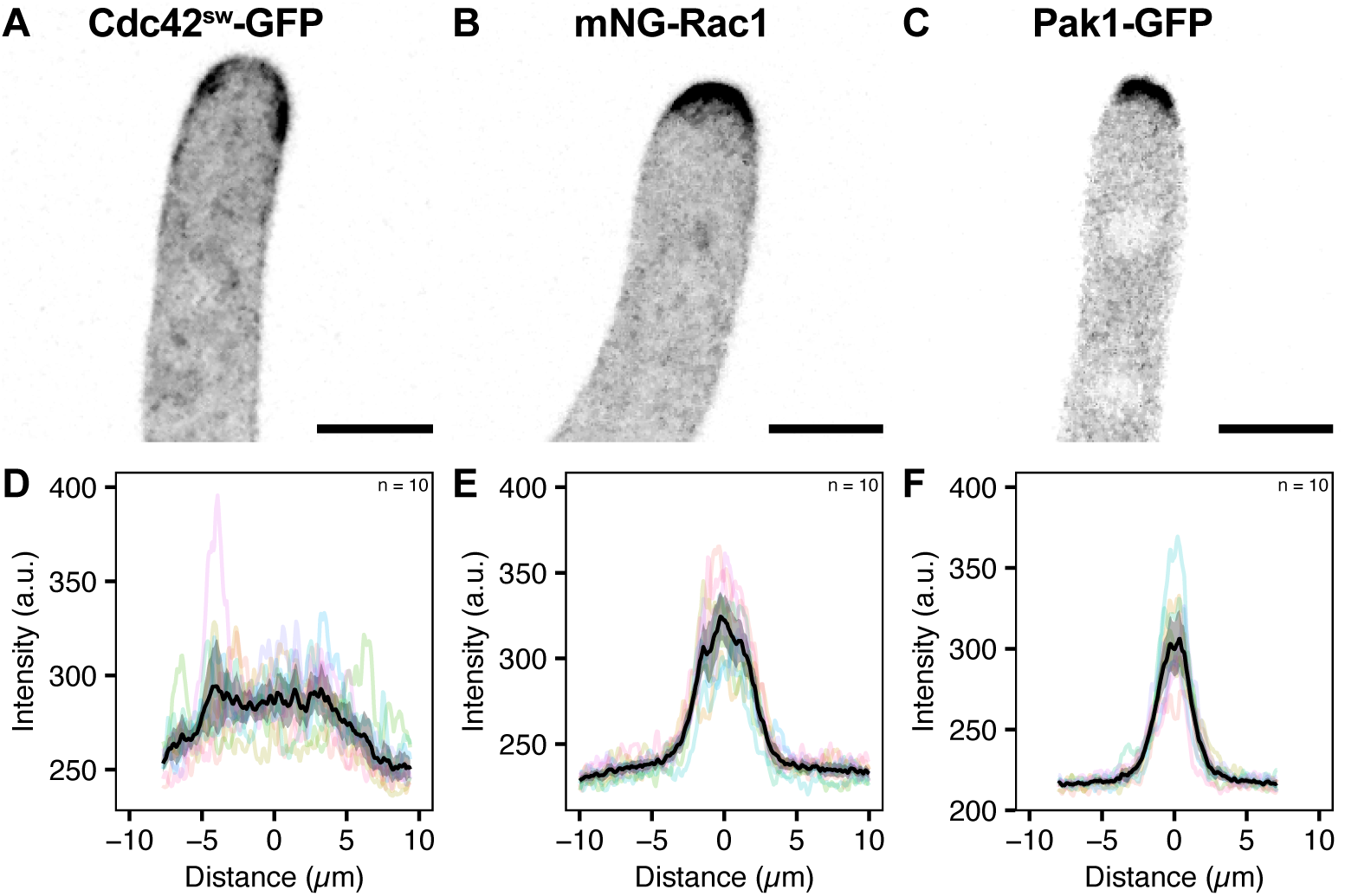
Localization of polarity factors in hyphae. (A-C) Maximum projection inverted confocal images showing localization of (A) Cdc42^sw^-GFP (DLY26533), (B) mNG-Rac1 (DLY26554), and (C) Pak1-GFP (DLY25103) at growing hyphal tips. Scale bars, 5 µm. (D-F) Quantification of polarity factor intensity at hyphal tips: (D) Cdc42^sw^-GFP (DLY26533) (E) mNG-Rac1 (DLY26554), (F) PAK1-GFP (DLY25103). Colored lines represent individual line scans performed on maximum projection images. Black line and shaded area indicate the mean intensity and 95% confidence interval for 10 hyphal tips in each case.

As expected, all three polarity factors localized to growing hyphal tips. Cdc42^SW^-GFP localized to the membrane in a broad crescent (Figure 1, A and D), whereas mNG-Rac1 (Figure 1, B and E) and Pak1-GFP (Figure 1, C and F) concentrated more tightly at growing hyphal tips. Polarity probes were concentrated at branch sites, septation sites, and budding sites (Supplemental Movie 2). Budding occurred from septated compartments and from tips that had stopped extending (Supplemental Movie 3).

### Characteristics of hyphae

As *A. pullulans* is an emerging model organism for fungal cell biology, we set out to characterize hyphal growth characteristics in this organism. Growing hyphae accumulated secretory vesicles marked by the vesicle Rab GFP-Sec4 at the tip (Figure 2A, Supplemental Movie 4). Using a fluorescent plasma membrane probe to outline the hyphae, we measured hyphal width at about 4 µm (Figure 2A-C, Table 1), wider than those of *Aspergillus nidulans* (3 µm) (Kaminskyj & Hamer, 1998) and *Candida albicans* (2 µm) (Sevilla & Odds, 1986), but narrower than *Neurospora crassa* hyphae (11.5 µm) (López-Franco et al., 1994). Hyphal branches emerged from the sides of the hyphae at irregular intervals (Figure 2E-F, Table 1). Hyphal growth rate was also somewhat variable. Tip growth was quantified by tracing the trajectory of Spa2-mCherry, a polarisome component that localizes to a clearly defined spot at growing hyphal tips with low cytoplasmic background (Figure 2G, Supplemental Movie 5) (Araujo-Palomares et al., 2009; Cao et al., 2021; Carbó & Pérez-Martín, 2008; Crampin et al., 2005; Knechtle et al., 2003; Meyer et al., 2008; Virag et al., 2007). Each track in Figure 2G represents a continuous period of hyphal extension. Hyphae grew in a persistent direction, with generally low tortuosity (Figure 2G-I). We noted several short tracks emerging initially, indicating short growth periods followed by pauses. These shorter tracks grew at variable speeds (Figure 2J and K), while longer tracks representing more mature hyphae extended processively at mean speeds of 15-20 µm/h (Figure 2K, Table 1). This extension rate is similar to that of *C. albicans* hyphae (∼ 15-20 µm/h) (Sevilla & Odds, 1986) but slower than those of *A. nidulans* hyphae (30 µm/h) (Horio & Oakley, 2005) and *N. crassa* hyphae (720 µm/h) (López-Franco et al., 1994). We also noted that mitosis occurs in a parasynchronous wave within hyphal compartments (Supplemental Movie 6). The wave did not spread to neighboring compartments separated by septa. Moreover, mitosis did not stop the extension of growing hyphal tips (Supplemental Movie 6). The metrics summarized in Table 1 provide a baseline quantitative characterization of hyphal growth in *A. pullulans* under our experimental conditions, although we note that growth patterns may vary in other conditions.

**Figure 2.**
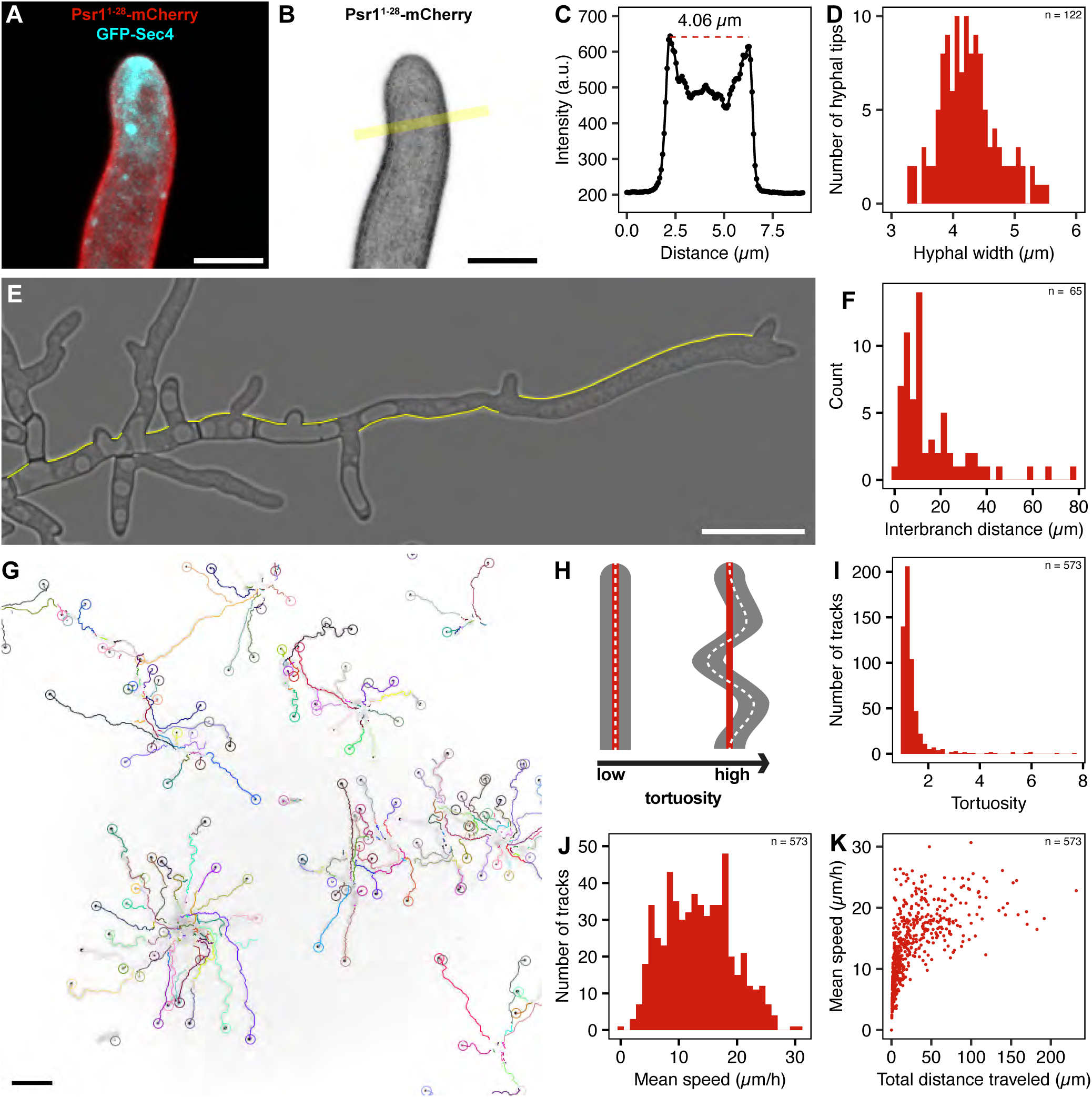
Hyphal growth characteristics. (A-C) Determining hyphal width. (A) Merged two-color maximum projection confocal image showing an example of a growing hyphal tip with GFP-Sec4 in cyan (vesicle marker accumulation at the tip indicates that this is a growing tip) and Psr1^1-28^-mCherry in red (plasma membrane marker) (DLY24564). (B) Inverted maximum projection of the same tip shown in A showing only Psr1^1-28^-mCherry channel with yellow line indicated to show location of the intensity measurement for the membrane probe. Scale bar, 5 µm. (C) Fluorescence intensity of Psr1^1-28^-mCherry signal along the yellow line shown in B. The dashed red line indicates the distance between plasma membrane peaks, yielding the width of the hypha shown in B. (D) Histogram of hyphal widths measured by the process shown in A-C (n= 122). (E) Example Differential Interference Contrast image of a hypha (DLY26075) with yellow lines illustrating how we measured the distances between branches along the hypha. Scale bar, 25 µm. (F) Histogram of interbranch distances measured as shown in E (n = 65). (G) Inverted fluorescence image of polarity marker Spa2-mCherry in hyphae (DLY25014) at the end of 17 h movie. Overlaid tracks followed Spa2-mCherry spots as hyphae grew. Tracks and circles indicated final tip location are colored by track ID. (H) Cartoon depicting low and high tortuosity hyphae. Tortuosity is calculated as the ratio between total distance traveled by a hypha (shown as dashed white line) and the net displacement (shown as red solid line). (I-K) Characteristics of hyphal growth tracks (n = 573) shown in G. (I) Histogram of tortuosity. (J) Histogram of mean speed. (K) Scatter plot showing relationship between mean speed and total distance traveled.

**Table 1:** Characteristics of *A. pullulans* hyphae.

|  | Mean | Median | SD | n |
| --- | --- | --- | --- | --- |
| Hyphal width | 4.3 $\mu\text{m}$ | 4.2 $\mu\text{m}$ | 0.5 $\mu\text{m}$ | 122 |
| Interbranch distance | 16.5 $\mu\text{m}$ | 10.8 $\mu\text{m}$ | 15.6 $\mu\text{m}$ | 65 |
| Track mean speed | 13.4 $\mu\text{m/h}$ | 13.2 $\mu\text{m/h}$ | 5.8 $\mu\text{m/h}$ | 573 |
| Tortuosity | 1.4 | 1.2 | 0.7 | 573 |

### Hyphal initiation

To understand how yeast cells switch to hyphal growth, we grew a strain harboring probes for nuclei (SpH2B-mCherry) and polarity sites (Pak1-GFP) under hyphal-inducing conditions. To our surprise, none of the cells present at the beginning of the movie directly initiated hyphae. Rather, they first proceeded through at least one and often several cell cycles that involved either budding cycles, or “nuclear cycles” in which the cells grew in size and doubled the number of nuclei without budding or undergoing cytokinesis (Figure 3A-C). We will refer to the cells present at the start of the movie as “F0” cells, and their daughter cells, born during the movie, as “F1” cells. After a variable number of budding and/or nuclear cycles, the cells polarized to form a protrusion that marked hyphal initiation (Figure 3A-C).

**Figure 3.**
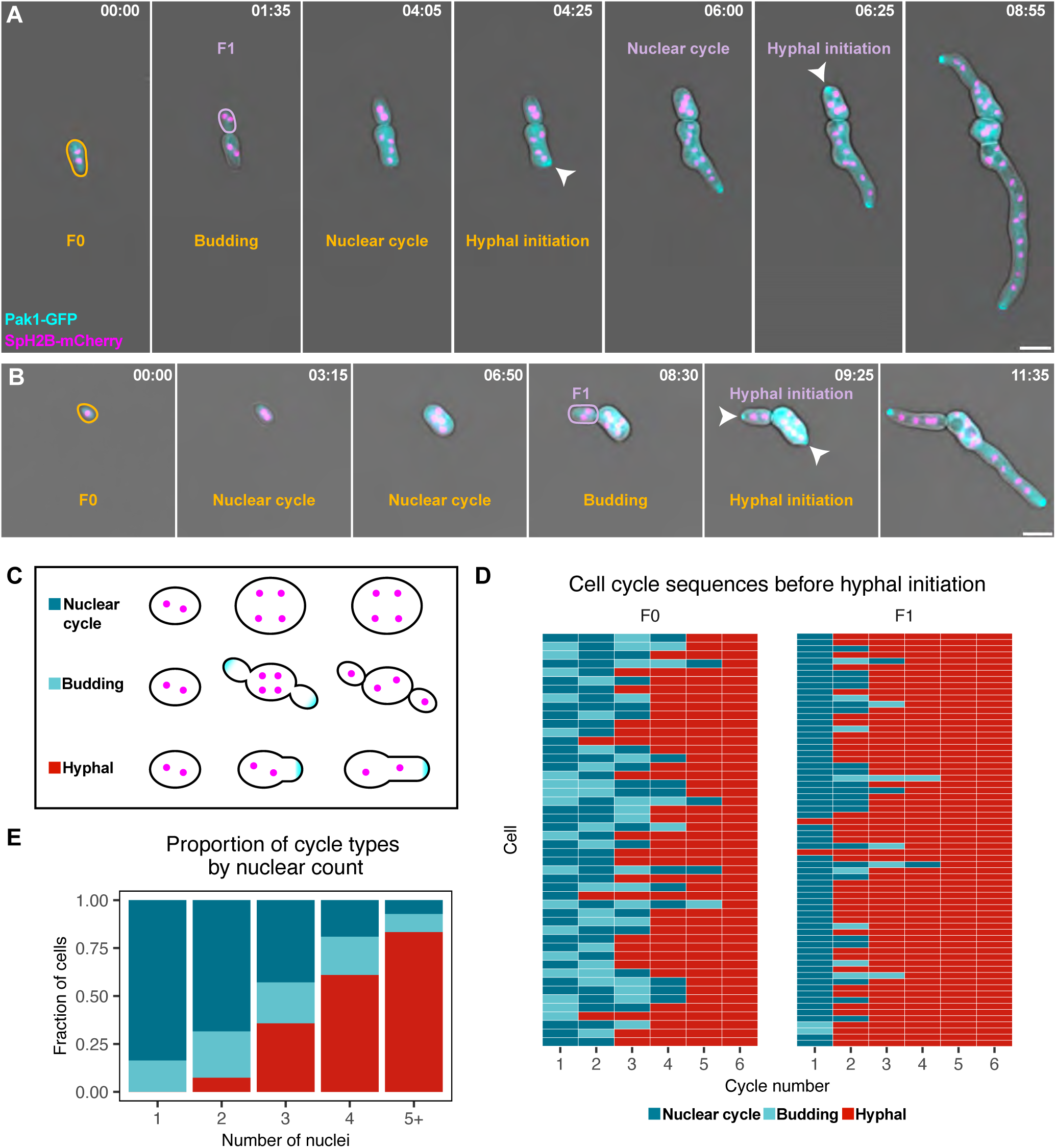
Hyphal initiation follows budding and nuclear cycles. (A-B) Two-color time-lapse imaging of nuclei (SpH2B-mCherry, magenta) and polarity sites (Pak1-GFP, cyan) in cells initiating hyphae (DLY25103). Differential Interference Contrast images merged with fluorescent channels. Cells present at the start of movie (F0 cells) are outlined in gold. Cells born during the movie from F0 parents (F1 cells) are outlined in light purple. Arrowheads point to hyphal initiation. Time shown in h: min. Scale bar, 10 µm. (C) Cartoon depicting nuclear cycle, budding cycle, and hyphal initiation. Nuclei depicted in magenta and polarity sites in cyan. (D) Tile plot illustrating cell cycles leading up to hyphal initiation in the same strain as in A-B. Each row represents a cell and columns represent the types of cell cycles that cell underwent. Nuclear cycles are shown in dark blue, budding cycles in light blue, and hyphal growth in red. F0 cells were present at the start of the movie (n = 48). F1 cells were born from F0 cells during the movie (n = 67). (E) Stacked bar chart showing proportion of cycle types undergone by same cells shown in D grouped by number of nuclei (n = 61, 136, 14, 105, and 42 for 1, 2, 3, 4, and 5+ nuclei, respectively). Nuclear cycles are shown in dark blue, budding cycles in light blue, and hyphal initiation in red.

There did not seem to be a stereotyped sequence or number of cell cycles needed before hyphal initiation. F0 cells (n = 48) showed considerable variation in number and type of cell cycles that occurred before hyphal initiation, while F1 cells (n = 67) more consistently underwent one or two nuclear cycles and then initiated hyphae (Figure 3A-D). In two instances, binucleate F1 cells initiated hyphae without first undergoing any nuclear or budding cycles (Figure 3B and D). Tracking the number of nuclei in F0 and F1 cells (n = 115) as they progressed to hyphal initiation (Figure 3E) indicated that the relationship between nuclear number and initiation was probabilistic, with the probability of cells initiating hyphae increasing with nuclear number (or possibly cell size, a correlated feature).

A large majority (86.5%, n = 104) of cells eventually initiated hyphae under our imaging conditions. However, most newly initiated hyphae did not grow continuously for very long. Of the cells that initiated, 75% (n = 88) paused growth within two hours. These young hyphae alternated between periods of growth and pausing (Figure 4A). The timing of pause onset after hyphal initiation (Figure 4B) and the duration of the pause (Figure 4C) were both quite variable. After the first pause, all cells reinitiated growth (n = 66), but only 74% reinitiated from the same site while the other 26% reinitiated growth from a different site. Some cells paused more than once, in a stop-and-start manner (Figure 4A).

**Figure 4.**
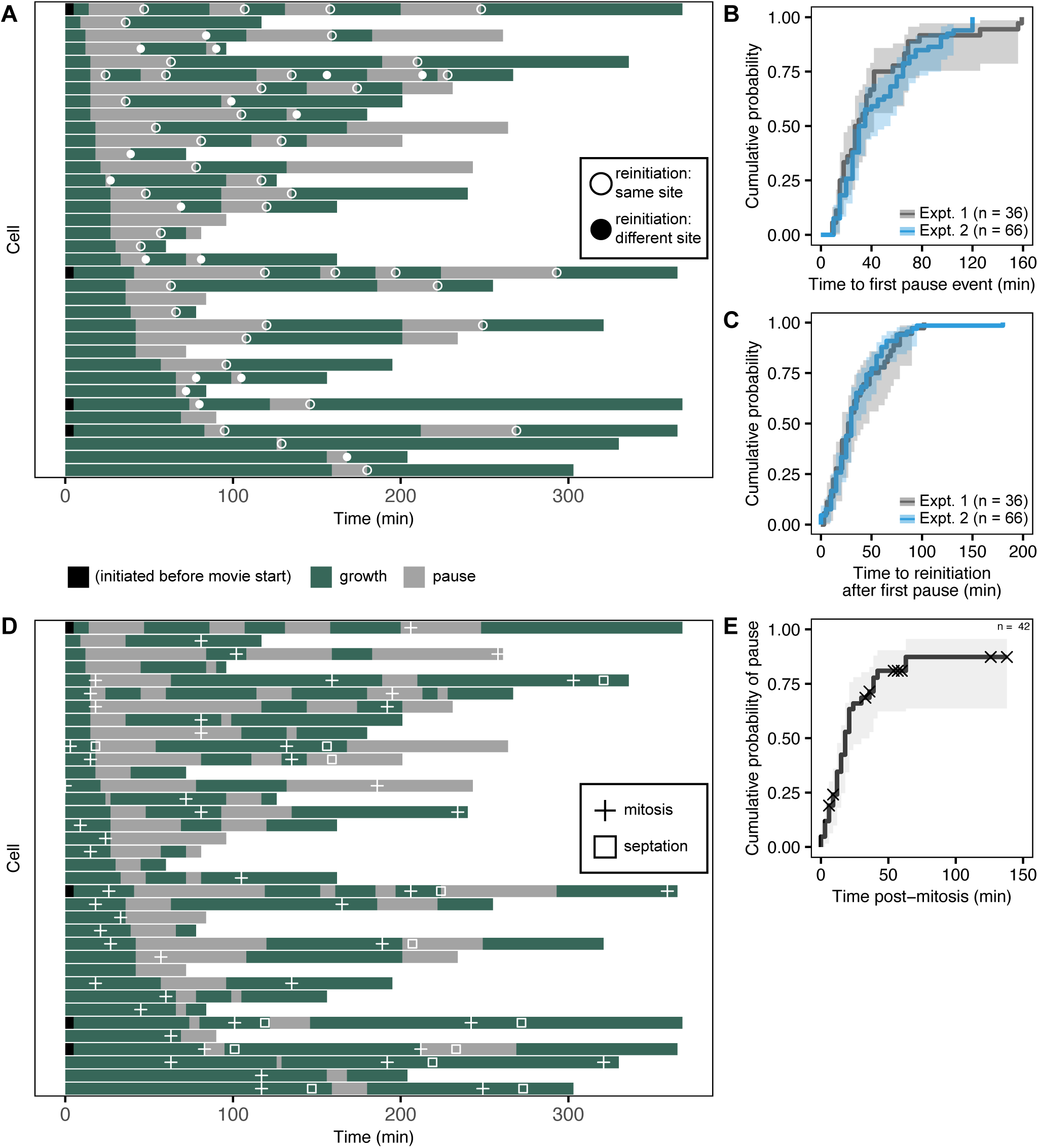
Initial hyphal growth is intermittent. (A) Time spent growing (green) or paused (grey) in newly initiated hyphae. Each row represents a cell with nuclei (SpH2B-mCherry) and polarity sites (Pak1-GFP) labeled (DLY25103) that was tracked by confocal time lapse imaging (n = 36 cells). Time zero represents yeast-to-hyphal transition (hyphal initiation) for each cell. Cells that were already polarized at the start of the movie are indicated by black bars. End time for each cell represents the end of the movie. Circles indicate whether resumption of hyphal growth after pausing occurred at the previous growth site (open circles) or elsewhere (filled circles). (B-C) Cumulative probability graphs showing (B) time to first pause event and (C) time to reinitiation of hyphal growth after first pause. Data collected from the same cells shown in A, grey line, n = 36) or from a separate experiment using widefield instead of confocal microscopy (DLY25014, blue line, n = 66). The shaded areas represent the 95% confidence interval for each experiment. (D) Same graph shown in A overlaid with symbols to indicate mitoses (+) and septation events (squares). (E) Cumulative probability of pause after mitosis event for mitosis events shown in D (n = 42). Censoring indicated by X symbols for cells where no pause was observed between mitosis event (time zero) and the end of the movie (X). The shaded area represents the 95% confidence interval.

Why would new hyphae stop and then restart growth? One possibility was that pausing was linked to the cell cycle—perhaps cells might depolarize during mitosis. However, although mitosis sometimes coincided with pausing, that was frequently not the case (Figure 4D). There were abundant instances of mitoses in growing hyphae with no pause, and conversely many pauses that were not correlated with any mitosis. The lack of a correlation between the onset of pausing and the time from the most recent mitosis (Figure 4E) suggests that pausing is not a cell cycle response. A similar lack of correlation was evident when comparing the timing of pauses and septation events (Figure 4D).

Examination of Pak1 localization revealed quite different patterns in cells that went through budding cycles, nuclear cycles, or hyphal growth. In budding cycles, mother cells developed small, focused patches of Pak1 at each bud site (Figure 5A, Supplemental Movie 7A). In nuclear cycles, cells failed to develop any localized Pak1 signal (Figure 5B, Supplemental Movie 7B). We also noted that cells sometimes developed transient Pak1-enriched zones that then disappeared (Figure 5C, Supplemental Movie 7C). In cells initiating hyphal growth, a single larger and more intense patch of Pak1 developed at the site of hyphal initiation (Figure 5D-E, Supplemental Movies 7C-D). When hyphae paused growth, Pak1 disappeared from the tip, and when they reinitiated growth Pak1 returned to the tip (or localized to a new patch when reinitiation occurred elsewhere) (Figure 5D-E). Thus, a multinucleate cell under the same growth conditions could develop a single large polarity site, several smaller polarity sites, or no stable polarity site.

**Figure 5.**
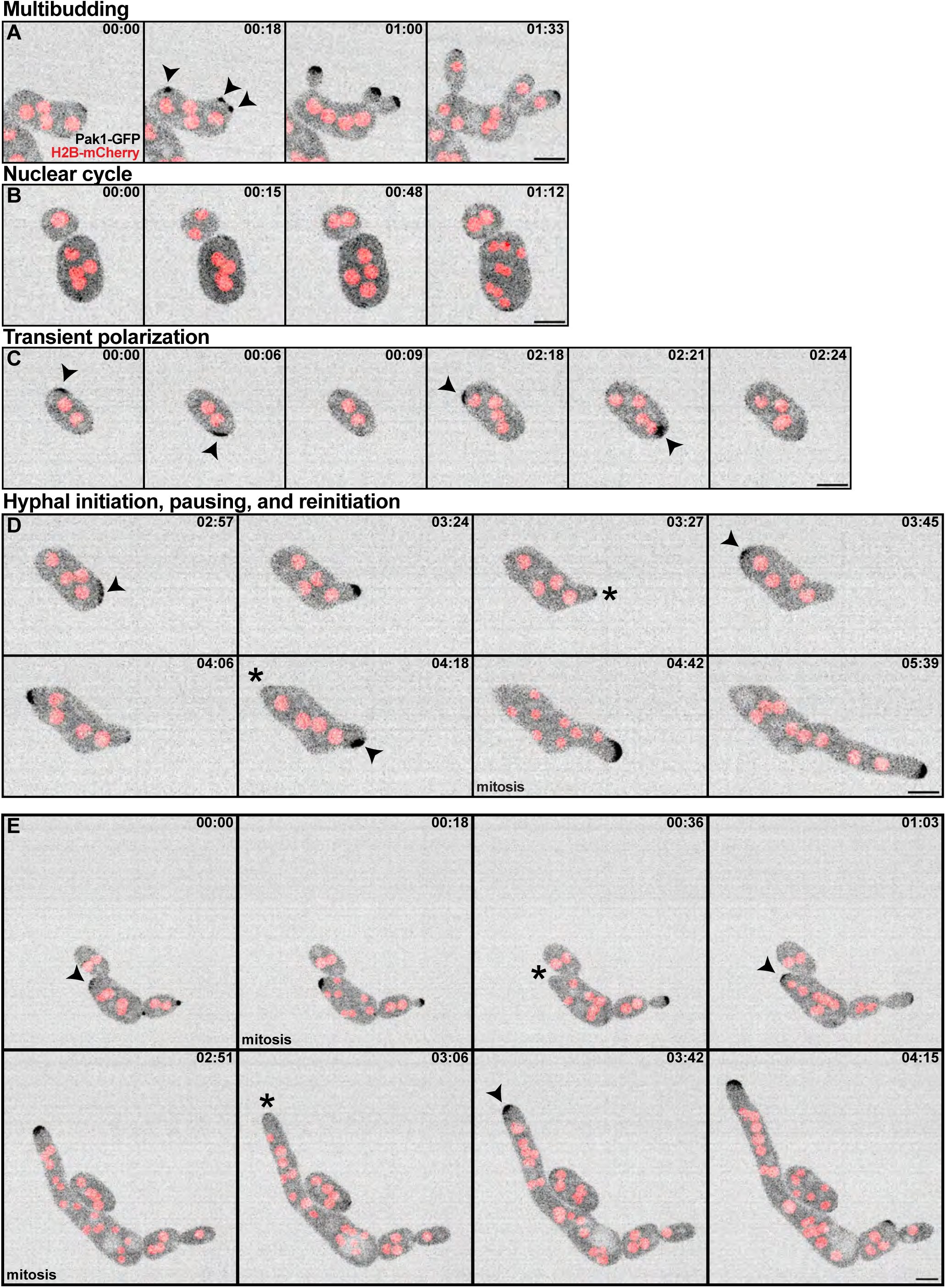
Pak1 localization during transition from yeast to hyphal growth. Maximum projection inverted confocal images showing localization of polarity sites (Pak1-GFP, black) and nuclei (SpH2B-mCherry, red) in growing cells before and after the switch from yeast to hyphal growth (DLY25103). (A) A multibudding cell generates multiple polarity sites. (B) Cells undergoing nuclear cycles do not show sustained polarization. (C) Transient polarization events not associated with polar growth. (D) Hyphal initiation, pausing, and reinitiation from a different site in the subsequent cell cycles of the same cell shown in C. (E) Hyphal initiation, pausing, and reinitiation from the same site. Arrowheads indicate new polarity sites. Asterisks indicate depolarization during pauses. Scale bars, 5 µm.

### Polarity requirements for hyphal growth

*A. pullulans* colonies contain a mixture of yeast and hyphal cells, with hyphae typically growing outwards at colony edges (Pollock et al., 1992). Hyphae were readily detected at edges of wild-type and *cdc42*Δ colonies (Figure 6, A and B), but not *rac1*Δ and *pak1*Δ colonies (Figure 6, C and D). To ask whether this failure was due to continued growth as yeast or to defective attempts to make hyphae, we imaged hyphal development. Wild-type and *cdc42*Δ cells formed hyphal networks (Figure 6, E and F), while *rac1*Δ and *pak1*Δ mutants formed shorter, wider structures whose tips sometimes appeared irregular and that had shorter compartments between septa (Figure 6, G and H). Thus, the failure to make hyphae was not due to continued yeast-form growth under these conditions. The nature of the defects is explored in more detail below.

**Figure 6.**
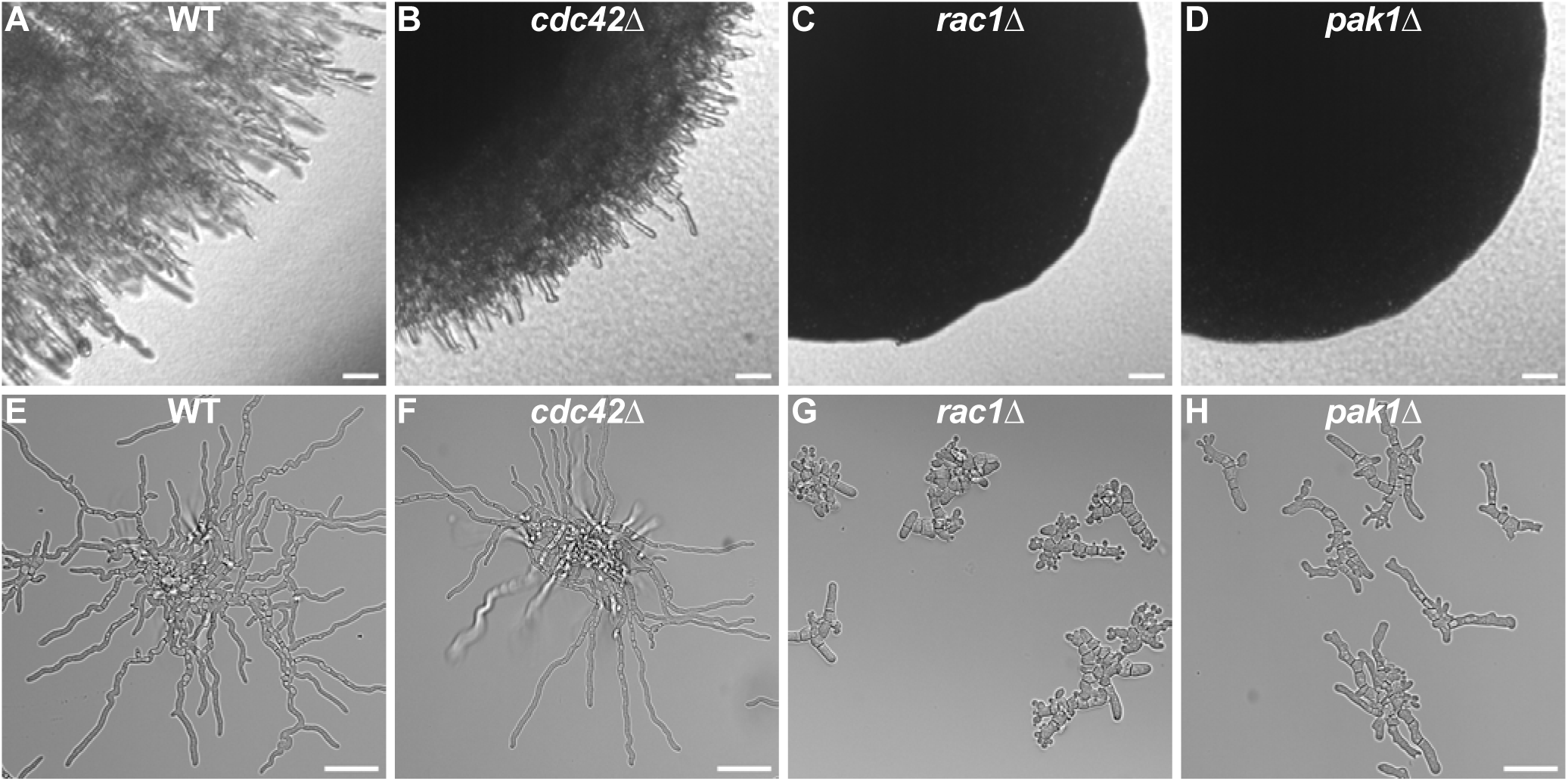
Rac1 and Pak1 are required for hyphal growth. (A-D) Brightfield images of colonies after 2 days of growth: (A) WT (DLY23540), (B) *cdc42*Δ (DLY25656), (C) *rac1*Δ (DLY25653), (D) *pak1*Δ (DLY25397). Scale bar, 50 µm. (E-H) Differential Interference Contrast images of cells grown under hyphal-inducing conditions for 18.5 h. Strains as in A-D. Scale bar, 50 µm.

### Hyphal initiation in mutants lacking the Pak1 kinase domain

It was not surprising that both *rac1*Δ and *pak1*Δ mutants had similar hyphal defects, as Pak1 appears to be a Rac1 effector (Crocker et al., 2025). To further investigate these morphology defects while still being able to track polarity, we used cells in which the kinase domain of endogenous Pak1 was replaced with GFP. We found that these cells had a similar mutant morphology to that of *pak1*Δ cells (Supplemental Figure S2). As shown previously (Wirshing et al., 2025), the Pak1 N-terminus (lacking the C-terminal kinase domain) still localizes to sites with active Rac1. Thus, using this Pak1ΔC-GFP probe, we were able to ask how Rac1 behaved in cells that were unable to make hyphae. We observed much broader zones enriched in Pak1ΔC-GFP than those observed with Pak1-GFP (Figure 7A-B), associated with slower extension of broader protrusions (Figure 7C). Thus, Pak1 kinase activity is needed to focus Rac1 activation at the hyphal tip.

**Figure 7.**
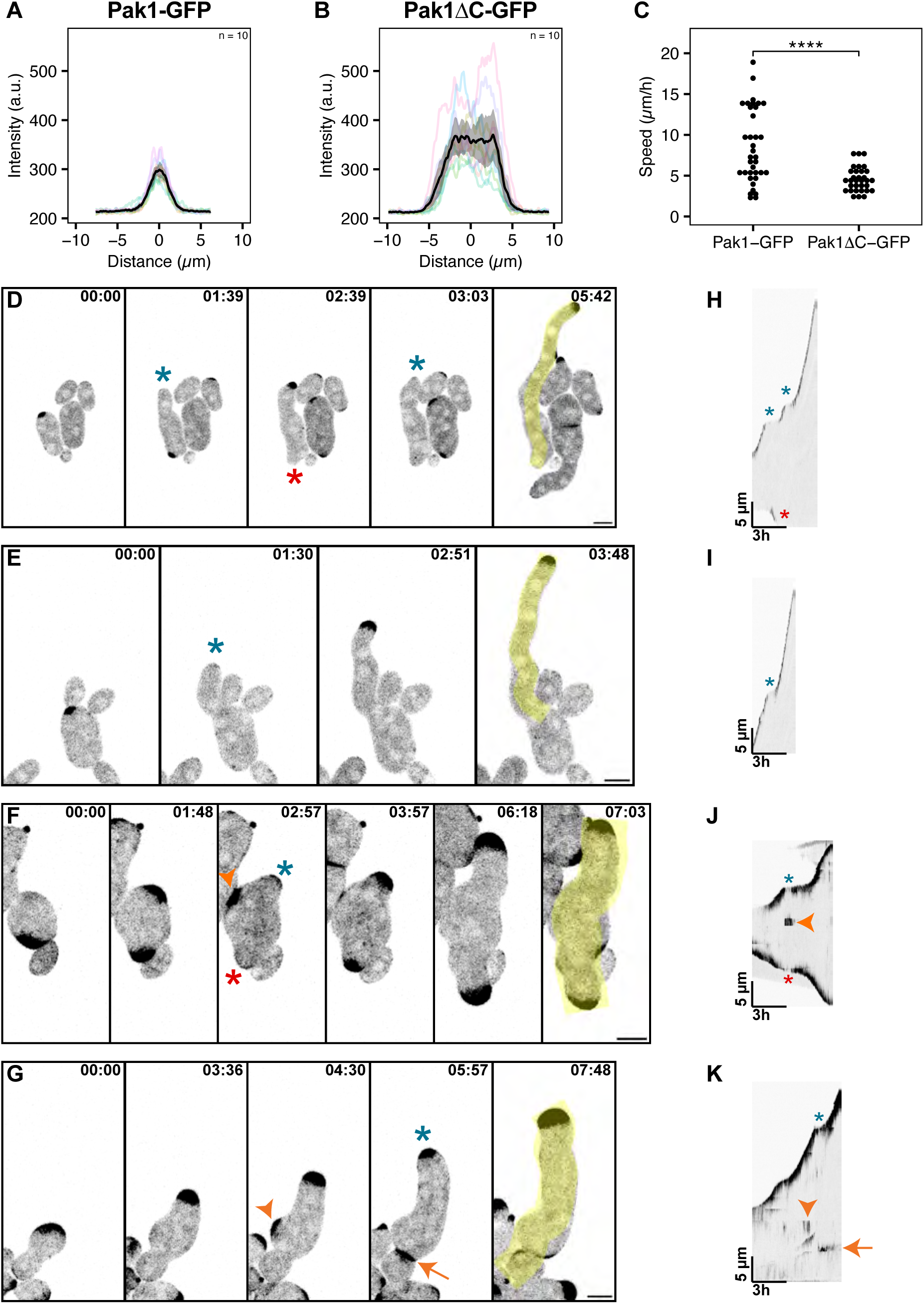
Mutants lacking the Pak1 kinase domain make multiple broad polarity sites. (A-B) Linescans of (A) Pak1-GFP (DLY25647) and (B) Pak1ΔC-GFP (DLY25650) analyzed as in Figure 1. N=10 hyphal tips. (C) Extension rate of polarized cells with Pak1-GFP (DLY25647, n = 36 growth periods across 9 tips) or Pak1ΔC-GFP (DLY25650, n = 32 growth periods across 9 tips). Asterisks represent statistical significance calculated from Welch t-test; p = 1.1 x 10^-5^. (D-G) Time-lapse maximum projection inverted confocal images showing localization of polarity sites in (D-E) WT cells (Pak1-GFP, black; DLY25647) and in (F-G) mutants lacking the Pak1 kinase domain (Pak1ΔC-GFP, black; DLY25650). Yellow lines on the last frame of each strip were used to create corresponding kymographs. Time shown in h:min. Scale bars, 5 µm. (H-K) Kymographs of cells shown in D-G, respectively. Asterisks label growth pauses (blue for upward-growing hyphae and red for downward-growing hyphae), arrowheads label additional polarity sites, and arrows label septation events.

The number, mobility, and persistence of polarity sites were also different in cells with Pak1ΔC-GFP (mutant) versus Pak1-GFP (wild-type). Early wild-type hyphae generally made a single polarity site at the hyphal tip, with only very transient bipolarity observed when switching the hyphal growth site (Figure 7D-E, H-U). In contrast, mutant cells frequently maintained co-existing broad polarity sites in different regions of a given cell, with gradual extension of all sites (Figure 7F, J, Supplemental Movie 8A-B). Wild-type polarity sites were stably positioned within the cell, whereas Pak1ΔC-GFP zones in mutant cells sometimes behaved like travelling waves (Supplemental Movie 8C-D). Like wild-type hyphae, the broad mutant projections extended in a stop-and-start manner, but whereas wild-type cells depolarized Pak1 during the pauses, mutant cells often retained polarized Pak1ΔC-GFP throughout the pause (Figure 7F-K).

Rarer but potentially informative behaviors were also observed. On some occasions, mutant cells appeared to undergo brief episodes of partial lysis, with cell cringing and brief depolarization followed by recovery (Supplemental Figure S3). In a few cases, mutant cells were observed to grow buds and broad protrusions simultaneously (Supplemental Movie 8A). The implications of these phenotypes for the role of Rac1 and Pak1 in hypha formation are discussed below.

## DISCUSSION

Here we describe the transition from yeast to hyphal growth in the polymorphic fungus, *A. pullulans*, with a focus on the regulation of cell polarity. Hyphal growth was broadly similar to that of related ascomycetes like *A. nidulans*, but the transition from yeast to hyphal growth modes displayed several unexpected features. These include (i) probabilistic switching between budding cycles, nuclear cycles, and hyphal initiation and (ii) stop-and-start growth of early hyphae. At the colony level, hyphal development required the polarity regulator Rac1 and its effector Pak1, but not Cdc42. At the cell level, our data suggest that *pak1*Δ mutant cells do attempt to construct hyphae but fail due to excessive broadening of the polarity site. As discussed below, the constellation of phenotypes exhibited by mutant cells raise the possibility that the *A. pullulans* polarity circuit is multi-stable and can switch between different behaviors based on small differences in initial conditions.

### Probabilistic hyphal initiation

Upon transfer to hyphal-inducing conditions, cells underwent a variable sequence of budding and nuclear cell cycles before initiating hyphal growth. Nuclear cycles involve DNA replication and mitosis but no bud formation or cytokinesis, yielding cells with approximately double the initial volume and double the initial number of nuclei. The probability of hyphal initiation increased as the cells accumulated more nuclei. We never observed hyphal initiation from cells with a single nucleus, but initiation frequency increased to >80% for cells with 5 or more nuclei. This means that for cells with intermediate numbers of nuclei (e.g. 4 nuclei), a cell may generate multiple small polarity sites to make multiple buds, or zero polarity sites to enter a nuclear cycle, or one larger polarity site to initiate a hypha. What factors influence the type of cell cycle and the appropriate number and size of polarity sites?

Morphological transitions in fungi are generally triggered by appropriate environmental stimuli, which can include nutrients, abiotic factors like temperature, host factors, pheromones, and quorum sensing factors. Such signals impact transcription regulatory networks, which are thought to impact the morphological growth program by regulating cell polarity, cell cycle progression, and cytoskeletal organization (Kijpornyongpan & Aime, 2021; Kurita et al., 2026; Nigg & Bernier, 2017). If hyphae and yeast forms represent distinct developmental pathways governed by transcription factor networks, then it could be that transcriptome differences between cells underlie the different behaviors of individual cells cultured under the same conditions. In this view, cells destined to undergo budding or nuclear cycles or hyphal initiation differ in their transcriptome in ways that predict their morphological development. These starting differences would be due to the particular life histories of each cell.

An alternative hypothesis to explain why different cells in the same culture embark on budding or nuclear cycles or hyphal initiation is that the polarity control circuit is multistable (Chiou et al., 2021). Control circuits that contain feedback loops, like the *A. pullulans* polarity circuit, can possess distinct “attractor” states (stable patterns of activity) and the attractor exhibited by a particular cell may depend on stochastic differences in the initial distribution of polarity proteins. In this view, the upstream transcription regulatory program need not differ among different cells in order to yield different outcomes. This possibility is consistent with the observation that mutants affecting the polarity regulator Pak1 can simultaneously engage in budding and hyphal extension, which are normally mutually exclusive outcomes.

### Early hyphal extension is unexpectedly episodic

Early hyphal growth occurred in a stop-and-start manner characterized by short bursts of tip growth interspersed with pauses. As hyphae grew longer, pauses became infrequent, and hyphae branched to make new tips from the sides of existing hyphae. As the hyphae matured, septation became more frequent, and some hyphal tips stopped growing. The non-growing tips and internal compartments then budded to produce new yeast-form cells.

Why would hyphae pause polar growth shortly after initiating? We investigated the possibility that pausing might be linked to the cell cycle. In *S. cerevisiae* and *S. pombe*, cells depolarize at specific cell cycle stages in a manner dependent on cyclin-dependent kinase activity (Chiou et al., 2017). However, depolarization was not tightly correlated with mitosis or septation in *A. pullulans*. In *S. pombe*, polarity proteins localize to the site of cytokinesis, where they function in septation (Gihana et al., 2021). Thus, it seemed plausible that septation might disrupt polarity at the growing tip by diverting polarity machinery to the site of septation. However, septation was not common during early hyphal growth of wild-type *A. pullulans*. In the Pak1ΔC-GFP mutants, we did observe septation events coinciding with paused growth and reduction of polarity at a nearby tip (Figure 7G, Supplemental Figure S4), suggesting that septation may be linked to pausing in this mutant, perhaps because of the unusual proximity of the septation site to the polarity site. On balance, however, septation does not explain the majority of pauses in young wild-type hyphae.

Another speculative possibility as to the cause of transient pausing is that hyphal volume expansion outpaces the rate at which protein synthesis can expand the amount of cytoplasm. In the constitutively filamentous *Ashbya gossypii*, the extension rate of young hyphae is slow, and gradually ramps up as the increasing volume of hyphal cytoplasm enables faster growth (Knechtle et al., 2003). In *A. pullulans* the growth rates of young hyphae were variable, but in some cases reached extension rates as fast as those of mature hyphae. Perhaps the pauses enable cytoplasmic content to “catch up” to a rapidly extending hypha. Another possible cause for pausing could be that initial hyphal extension creates cell wall vulnerabilities that need to be repaired before hyphal growth can continue safely (Municio-Diaz et al., 2022). Whatever the impetus that causes tip extension to pause, it appears to affect only young hyphae, as hyphal growth becomes continuous in longer hyphae.

### Role of Rac1 and Pak1 in hyphal morphogenesis

In *A. pullulans*, both *cdc42*Δ and *rac1*Δ mutants are capable of polarizing to form buds, indicating that either GTPase is sufficient to polarize (Crocker et al., 2025). Here, we report that under hyphal-inducing conditions, *rac1*Δ and *pak1*Δ mutants do not form normal hyphal networks. This was not due to a failure to polarize; instead, it was associated with overly broad and numerous polarity sites, which failed to focus growth to a single narrow tip. Pak1 is thought to be a Rac1 effector (Crocker et al., 2025). The similar phenotypes of both mutants indicate that Pak1 is likely the main Rac1 effector for hyphal development. The exact role of Pak1 in hyphal morphogenesis remains unclear, but Pak1 has been implicated in negative feedback during budding (Crocker et al., 2025). It seems possible that the overly broad and numerous polarity sites are a result of a lack of negative regulation by Pak1.

Wild-type *A. pullulans* cells display multiple polarity sites during multibudding and during branching of more mature hyphae. However, initial hyphal extension is characterized by a single polarity site at the hyphal tip. Young hyphal cells may transiently have two polarity sites (especially as they switch between growing from one side of the cell to another), but one is quickly lost while the other is maintained. This is reminiscent of competition between polarity sites in the single-budding yeast *S. cerevisiae* that guarantees cells only maintain one polarity site (Wu et al., 2015). Thus, despite having the ability to initiate multiple sites, something about the hyphal growth program in *A. pullulans* turns off additional polarity sites. This negative regulation likely involves Pak1 activity as mutants lacking the Pak1 kinase domain frequently maintain multiple polarity sites. In the *rac1*Δ*, pak1*Δ, and Pak1ΔC-GFP mutants, septation and budding from hyphae were observed earlier than in wild-type development. Wild-type hyphae usually septate and produce yeast cells once they are more mature. Thus, disruption of negative feedback may also be leading to premature septation and budding.

As with *A. pullulans*, filamentation in the dimorphic plant pathogen *Ustilago maydis* requires Rac1 but not Cdc42 (Mahlert et al., 2006). This very distantly related basidiomycete makes very thin dikaryotic filaments upon mating, revealing a very different cell biology than that of filamenting *A. pullulans* (Vollmeister et al., 2012). Interestingly, however, *U. maydis* filamentation also involves a PAK-mediated negative feedback loop that degrades the Rac1 exchange factor (Frieser et al., 2011). Short-circuiting the negative feedback by stabilizing the exchange factor results in cells that grow shorter, broader filaments (Frieser et al., 2011).Thus, despite their evolutionary divergence and distinct filamentation biology, both *U. maydis* and *A. pullulans* may use similar strategies to focus polarity to the hyphal tip. Further investigation is needed to uncover the mechanisms of polarity regulation in *A. pullulans*.

## MATERIALS AND METHODS

### *A. pullulans* strain construction and maintenance

All *A. pullulans* strains used in this study (Supplemental Table 1) were generated in the *EXF-150* strain background (Gostinčar et al., 2019) as described below. Strains were cultured in YPD (2% glucose, 2% peptone, 1% yeast extract) plates with 2% BD Bacto^TM^ agar (214050, VWR) at 24°C.

As tagging Cdc42 at its C-or N-terminus has been reported to interfere with localization and function, a sandwich tag was used to internally tag this protein (*CDC42-GFP^SW^*) (Bendezú et al., 2015; Woods et al., 2015). To replace the endogenous *CDC42* and *RAC1* with internally-tagged *CDC42* and N-terminally tagged *RAC1*, respectively, we used a pop-in/pop-out strategy. In the first (pop-in) step, a *ura3* auxotrophic strain (DLY24148) was transformed with linearized plasmid carrying the relevant tag as well as an intact *URA3* gene, digested at a unique site to target integration at the endogenous *CDC42* or *RAC1* locus. Transformants were selected for uracil prototrophy followed by colony PCR screening to confirm successful integration. These strains contain a locus with adjacent tagged and untagged versions of *CDC42* or *RAC1*, separated by plasmid sequences including *URA3*. In the second (pop-out) step, spontaneous recombinants that delete the intervening *URA3* were selected using 5-FOA (Boeke et al., 1984). Briefly, strains were grown in YPD+2% glucose overnight and ∼5×10^5^ cells were plated on CSM+2% glucose+10% YPD+1g/L 5-FOA. 5-FOA is lethal to cells containing an intact *URA3* gene. 5-FOA-resistant “pop-out” strains delete one copy of the GTPase as well as *URA3*, and further screening by colony PCR was used to verify retention of the fluorescently tagged loci.

The resulting locus was PCR-amplified and sequenced using Nanopore (Plasmidsaurus) to confirm sequence integrity and scarless replacement of the untagged gene with the tagged version.

Deletions lacking Cdc42 (protein ID 330431) and Rac1 (protein ID 341125) were constructed by replacing the open reading frame with a hygromycin resistance cassette using the 3-fragment PCR method previously described (Colarusso et al., 2025). To create a strain with a fluorescently labeled plasma membrane, we codon optimized the first 28 amino acids of *S. cerevisiae* Psr1 for *A. pullulans* and fused it to a codon optimized mCherry. This probe undergoes myristoylation and palmitoylation, which attaches it to the plasma membrane (Kuo et al., 2014; Siniossoglou et al., 2000). A single plasmid, DLB4784, was constructed to express Psr1^1-28^-mCherry, driven by the Chytrid *Spizellomyces punctatus* histone H2A promoter, and the exocytic vesicle marker GFP-Sec4, driven by the *scACT1* promoter. As shown previously, Sec4 (protein ID 271553) localizes to sites of active growth where exocytosis occurs (Wirshing et al., 2025). The Psr1^1-28^-mCherry and GFP-Sec4 cassettes were cloned into a backbone containing the *URA3* coding sequence in between holomogy arms that allow for integration at the *<u>URA3</u>* <u>locus using the</u> NEBuilder HiFi DNA Assembly Master Mix (E2621L, New England Biolabs) (Wirshing et al., 2024). DLB4784 was digested with ApaI and NruI-HF, releasing a linear fragment that was transformed into a *ura3::HYG_R_* auxotrophic strain (DLY24148) via PEG-mediated transfer. Transformants were validated by screening for URA prototrophy and hygromycin sensitivity.

Tagging of native SPA2 homolog (protein ID 274823) with mCherry was done using the 3-fragment PCR method previously described (Colarusso et al., 2025).

### *A. pullulans* hyphal induction & live cell microscopy

Low cell densities promote hyphal growth while higher densities increase budding (Finlay, 1987; Goshima, 2022; D. Park, 1984; Ramos & García Acha, 1975). In addition, we found that pre-culturing cells without agitation improved the fraction of cells that formed hyphae.

Cells were streaked out from frozen stocks onto YPD agar plates and grown at 24°C for 2-3 days. A single colony was inoculated into 5 mL of complete synthetic media (2% glucose, 6.71 g/L BD DifcoTM Yeast Nitrogen Base without Amino Acids, BD291940, FisherScientific, and 0.79 g/L Complete Supplement Mixture, 1001-010, Sunrise Science Products). Cultures were inoculated from this stock at fivefold serial dilutions and incubated without agitation at 24°C for ∼24 hours. Cultures that had reached a low density of 1.0-2.2 x 10^6^ cells/mL were selected. If multiple strains were being compared for their ability to form hyphae, culture density was normalized at this step. Then, 2 µL of culture was placed into an 8-well glass-bottomed Ibidi chamber (80827, Ibidi) and covered with a pad of CSM with 5% agarose (97062-250, VWR). 5% agarose was necessary to decrease the frequency of hyphal tips burrowing into the agarose pad and growing out of the focal plane of the microscope (Supplemental Figure S5). The ibidi chamber was either imaged directly or incubated at 24°C in a humidity chamber (an empty pipette tip box with a layer of water) before imaging as described below. All microscopy was conducted at room temperature (20-22°C).

To capture the entire process from hyphal initiation to network formation, chambers were mounted on the microscope and imaged every 5 min for 17-24 h (Figures 3G-K, 4, and 5B-C; Supplemental Movies 1 and 5). These movies were captured on a widefield Nikon ECLIPSE Ti2 inverted microscope with a Plan Apo 20× air objective (NA 0.75; Nikon Instruments), an sCMOS pco.edge camera (Excelitas Technologies), and an X-Cite XYLIS LED Illumination System (Excelitas Technologies) controlled by NIS-Elements software (Nikon Instruments). A single medial plane was imaged, using 50-100 ms exposure times at 5-20% laser power (488 nm and 561 nm). To determine the number of nuclei required for hyphal initiation, images were acquired at 7 z-slices (1 µm step intervals) in order to capture all nuclei.

To image the early steps in hyphal initiation, cells were incubated in the ibidi chamber for ∼ 5h before imaging. Movies were captured on a Nikon Ti2E inverted confocal microscope with a CSU-W1 spinning-disk head (Yokogawa), a CFI60 Plan Apochromat Lambda D 60× Oil Immersion Objective (NA 1.42; Nikon Instruments), and a Hamamatsu ORCA Quest qCMOS camera controlled by NIS-Elements software (Nikon Instruments). Images were acquired at 19 z-slices (0.6 µm step intervals) every 3 min for 6-12h. The SpH2B-mCherry probe was imaged with a 50 ms exposure time at 5% laser power (561 nm). The Pak1-GFP and Pak1ΔC-GFP probes were imaged with a 150 ms exposure time at 4.5-12% laser power (488 nm).

To image mature hyphae, cells were left to incubate in the ibidi chamber overnight and photographed the next day (within 14-24h). Differential Interference Contrast images of hyphae were captured at a medial plane with the widefield microscope setup described above (Figure 6E-H; 2E). Images of strains with tagged polarity proteins (Figure 1A-C, Supplemental Movies 2-3, Supplemental Figure 2) were captured with the confocal microscope setup described above. Images were acquired at 19 z-slices (0.6 µm step intervals). The SpH2B-mCherry probe was imaged with a 50 ms exposure time at 5% laser power (561 nm). The polarity probes were imaged with a 150 ms exposure time at 15-30% laser power (488 nm). Imaging to validate the Psr1^1-28^-mCherry GFP-Sec4 strain (Supplemental Movie 4) was acquired with a 200 ms exposure time at 10% laser power (488 and 561 nm). Images taken to determine hyphal width were captured at a wider z-range to ensure hyphae of various sizes were captured (Figure 2A-B). Images were acquired at 27 z-slices (0.6 µm step intervals) with a 200 ms exposure time at 20% laser power (488 nm and 561 nm). Movies of mitosis with SpH2B-mCherry and 3xGFP-Tub2 probes (Supplemental Movie 6) were acquired at 15 z-slices (0.6 µm step intervals) with a 175 ms exposure time and 5% laser power (488 nm and 561 nm). One movie was imaged every 30 seconds for 15 minutes and another was imaged every 2 minutes for an hour. Images of colonies were taken on a Singer Instruments MSM 400 dissection microscope with the 4x objective after colonies were grown on YPD plates for 2 nights at 24°C (Figure 6A-D).

### Image analysis

The median filter in NIS-Elements General Analysis 3 (GA3, Nikon Instruments) was used to denoise confocal images and movies. Maximum intensity projections were created for files with multiple z-slices.

The NIS.ai software’s Denoise.ai filter was used to denoise fluorescent widefield images and movies. When necessary (Figure 3A-B and Supplemental Movie 1), the DIC channel was bleach corrected using a simple ratio with background intensity set to zero (Miura, 2020)

Images that had only brightfield or DIC but no fluorescent channels (Figures 2, 3E, Supplemental Figure S5) were not denoised.

### Measuring spread of polarity factors at hyphal tips

The spread of polarity probes at hyphal tips was measured by tracing a line scan along the hyphal tip using the segmented line tool in FIJI, set to 10 pixels in width. This process was done for 10 representative hyphal tips per probe and lines were saved as ROIs. Hyphal tips that were growing straight were selected (not actively turning or budding). For each image, all line ROIs were selected, plotted using “multiplot” within the ROI manager, and the corresponding data was exported for analysis. Intensity along those line scans was plotted with each line being centered to the center of the hyphal tip.

### Measuring hyphal width

Hyphal width was measured using a strain with a fluorescently-labelled plasma membrane (DLY24564). Using the straight line tool in FIJI, a line was drawn 5 µm back from hyphal tips, orthogonal to the growth direction of the hypha (Figure 2A).

In our images, 5 µm was determined in FIJI to be ∼65 pixels. In order to place the line 5 µm back from the tip, line width was set to 130 pixels (such that there are 65 pixels of shaded area on either side of the line) and the line was positioned so that the edge of the shaded area touched the edge of the hyphal tip. Then, the line width was adjusted to 10 pixels and saved as an ROI (Figure 2B). This process was repeated to analyze a total of 122 actively growing hyphal tips. Hyphal tips were considered to be actively growing if they showed an accumulation of GFP-Sec4 signal at the tip, indicating vesicle secretion directed at the tip (Supplemental Movie 4).

Hyphal tips were excluded if they were not actively growing (no GFP-Sec4 signal), were growing out of focus (determined by looking at brightfield), were budding, were shorter than 20 µm, or if they were bent/pressed up against another membrane.

Width of each hyphal tip was determined by plotting all intensity values along its corresponding ROI line. Since the signal should be highest at the edges of the cell, intensity data was fit to a spline and the two highest peaks were identified (corresponding to the two edges of the hypha). The distance between those two peaks was reported as the hyphal width (Figure 2C).

### Measuring interbranch distance

Interbranch distance was measured by tracing the distance between the end of one branch and the start of the next adjacent branch using the segmented line tool in FIJI (Figure 2E). Interbranch distance was only measured between primary branches, but not secondary branches (branches that emerge on primary branches).

### Measuring tortuosity and speed of hyphal extension

TrackMate was used to obtain tracks corresponding to hyphal extension during ∼ 17h of hyphal growth of DLY25014 (Ershov et al., 2022; Tinevez et al., 2017). The frames included in the tracking were from the first time frame until the frame where hyphal tips started growing out of focus. To detect Spa2-mCherry spots, we used the Laplacian of Gaussian detector with an estimated object diameter of 3.5, a quality threshold of 1.7, and subpixel localization. Tracks were created with the Kalman tracker using an initial search radius of 3 µm, a search radius of 3 µm, and a max frame gap of 1. This frame gap was included since the movie was captured at a medial plane and the hypha may have grown out of focus for a frame and returned at the next one.

Tracks were then filtered to keep only tracks with a mean quality threshold above 1.98, more than 2 spots in track (2-spot tracks often represented cytokinetic events, or transient polarization that does not have associated growth), and a confinement ratio above 0.13 (low confinement ratio tracks often represented debris). After this initial filtering, additional tracks were manually removed if they (1) did not represent true growth signal, (2) tracked bud growth, or (3) grew out of focus as determined by DIC channel.

The TackMate output provided track mean speed and total distance travelled by tracks. Tortuosity was calculated for each track by dividing total distance traveled by straight-line distance between start and end points. This analysis was conducted for 2 stage positions, resulting in 573 total tracks after filtering.

### Tracking cell cycles leading up to hyphal initiation

To track cell cycles leading up to hyphal initiation, we imaged cells with SpH2B-mCherry (nuclei) and Pak1-GFP (DLY25103). Cells present at the start of the movie were designated “F0” cells and cells born during the movie from F0 cells were designated “F1” cells. All F0 and F1 cells that initiated hyphae were labelled and tracked to determine the number and kind of cell cycles they underwent before hyphal initiation. The number of nuclei at the beginning of the movie or at birth, and in each subsequent cell cycle leading up to hyphal initiation was recorded. Initiation was defined as the first time point in which a cell had localized signal of Pak1-GFP at a protrusion (that went on to grow in a polar manner).

### Tacking hyphal initiation, pausing, and re-initiation

Pausing and reinitiation were tracked initially in the widefield microscopy movies (images taken every 5 min) of DLY25014 with overlaid Spa2-mCherry tracks described above (Figure 4B-C). For each F0 and F1 cell, we noted whether the cell initiated a hypha during the movie, the time point at which it initiated, if it paused growth (pause in apical extension and loss of Spa2-mCherry signal), the time point at which it paused (first time point where Spa2-mCherry signal is lost), if it reinitiated growth, the time point at which it reinitiated, and whether reinitiation was from the same site that was previously growing or a different site. Spa2-mCherry tracks were used as guides but timing of growth (re)initiation and pauses was manually determined for each cell. The duration of the first growth and first pause was calculated by taking the difference between the time of (re)initiation and pause. This analysis was conducted for 2 stage positions, with a total of 104 cells.

A similar analysis was performed on confocal movies (images taken every 3 min) of DLY25103 with nuclear (SpH2B-mCherry) and polarity (Pak1-GFP) probes (Figure 4) to track periods of growth and pausing. The higher time resolution of these movies also allowed for tracking of mitosis, which lasts ∼5 minutes in *A. pullulans* (Petrucco et al., 2025). Mitosis and septation were manually annotated. Mitosis was called at the first frame in the movie where the nuclei separated. Septation was called at the first frame where the septum became visible in the brightfield channel.

### Generating kymographs and measuring hyphal growth speed

Confocal movies of PAK1-GFP (DLY25647) and Pak1ΔC-GFP (DLY25650) strains were cropped to capture initial polar growth of cells of interest. Images were rotated so that the hypha grows upwards for the first growth period. The segmented line tool in FIJI was used to draw a line from the back of the cell to the growing tip (or from bottom to top) at the last time frame (Figure 7D-G, Supplemental Figure 4A-C). Linewidth was adjusted to capture the width of the hyphae (50 pixels for WT DLY25647, 100 pixels for wider DLY25650 mutants). Each line was saved as an ROI and KymographBuilder plugin in FIJI was used to generate kymographs (Mary et al., 2016). Kymographs were rotated 90° to the left so that time is on the x-axis and the greyscale image was inverted to match the corresponding time lapse images.

To determine speed of early growth of these strains (Figure 7C), the segmented line tool (linewidth 10 pixels) was used to trace polarity signals along the kymographs for both strains. A flat line segment was drawn on portions of the kymograph that corresponded to periods where the hyphae stopped growing. A total of 9 hyphal tips were traced for each strain. Lines were saved as ROIs and their coordinates were exported. Slopes of each line segment were calculated to determine speed. Periods of hyphal growth pausing were filtered out by excluding segments that had speeds below 2 µm/h. This resulted in a total of 36 growth periods for DLY25647 and 32 growth periods for DLY25650.

### Growth assay

To compare growth of different strains, a single colony of each strain was inoculated into 5 ml of YPD (2% glucose) and incubated on a rotating wheel at 24°C for 24 hours. Cultures were then diluted to an OD_600_ of 2 and serially diluted (10-fold dilutions) 5 times in sterile deionized water in a 96-well plate. Dilutions were transferred to YPD plates with a 48-pin metal microplate replicator. Plates were grown at 24°C or 30°C and imaged on a scanner (Epson Perfection V600 Photo) after 24-48 hours. Two replicate plates were prepared for each temperature.

## ACKNOWLEDGEMENTS

Thank you to members of the Lew lab for stimulating discussion and feedback on this manuscript. Thank you to Analeigha V. Colarusso for DLY25014 strain construction. Thank you to Erin R. Curtis for coding in R expertise. This work was funded by NIH/NIGMS grant R35GM122488 to DJL.

