## supplement for "Regulation of polarity during morphological transitions in *Aureobasidium pullulans*"

**TABLE S1.** ***Aureobasidium*** **strains used in this study**

All strains are in the EXF-150 strain background.

| Strain | Genotype | Source |
|---|---|---|
| DLY23540 | WT EXF-150 | Gostinčar et al., 2014 |
| DLY24148 | *ura3::HYG$^{R}$* | Wirshing et al., 2024 |
| DLY24564 | *URA3:PSR1$^{1\text{-}28}$-mCherry:GFP-SEC4* | This study |
| DLY24568 | *URA3:SpH2B-mCherry:3xGFP-TUB2* | Petrucco et al., 2025 |
| DLY25014 | *SPA2-mCherry:NATR, URA3:3xGFP-TUB2* | This study |
| DLY25103 / CPY519 | *PAK1-GFP:HYG$^{R}$, URA3:SpH2B-mCherry* | Crocker et al., 2025 |
| DLY25397 | *pak1::HYG$^{R}$* | Cocker et al., 2025 |
| DLY25647 | *URA3:spH2Bp-3xmCherry; PAK1-GFP:NAT$^{R}$* | Wirshing et al., 2025 |
| DLY25650 | *URA3:spH2Bp-3xmCherry; pak1ΔC-GFP:NAT$^{R}$* | Wirshing et al., 2025 |
| DLY25653 | *rac1::HYG$^{R}$* | This study |
| DLY25656 | *cdc42::HYG$^{R}$* | This study |
| DLY26075 | *URA3 : SpH2B-mCherry : mNeonGreen-TPM1* | Colarusso et al., 2026 |
| DLY26533 | *CDC42$^{sw}$-GFP, URA3:SpH2B-mCherry* | This study |
| DLY26554 | *mNeon-RAC1, URA3:SpH2B-mCherry* | This study |

### SUPPLEMENTAL FIGURE LEGENDS

**Supplemental Figure 1. Colony growth and hypha formation of A. pullulans strains used in this study.** (A) YPD plates with 10-fold serial dilutions of cultures of the indicated strains were grown for the indicated times at the indicated temperatures. Strains from top to bottom in the top plate: DLY23540, DLY26533, DLY26554, DLY25103, DLY25656, DLY25653, and DLY25397 and the bottom plate: DLY23540, DLY24564, DLY26075, DLY25014, DLY25647, and DLY25650. (B-D) Brightfield images of colonies after 2 days of growth: (B) Cdc42$^{sw}$-GFP (DLY26533), (C) mNG-Rac1 (DLY26554), and (D) Pak1-GFP (DLY25103) Scale bar, 50 µm.

**Supplemental Figure 2. Effect of deleting the Pak1 kinase domain on hyphal development.** Brightfield images of (A) Pak1-GFP (DLY25647) and (B) Pak1ΔC-GFP (DLY25650) mutant cells after 21.5 h of growth under hyphal-inducing conditions. Scale bar, 25 µm.

**Supplemental Figure 3. Brief depolarization events in Pak1ΔC mutants.** Time-lapse inverted fluorescence imaging (top) and Differential Interference Contrast imaging (bottom) of Pak1ΔC-GFP (DLY25650) cells. Asterisk labels transient depolarization and arrow points to vacuoles. Scale bar, 5 µm.

**Supplemental Figure 4. Septation sometimes coincides with growth pausing in Pak1ΔC-GFP mutants.** (A-C) Time lapse maximum projection inverted confocal images showing localization of polarity sites (Pak1ΔC-GFP, black) in mutants lacking the Pak1 kinase domain (DLY25650). Yellow lines were used to create kymographs. Time shown in h:min. Scale bar, 5 µm. (D-F) Kymographs of cells shown in A-C, respectively. Asterisks label growth pauses, arrowheads label additional polarity sites, and arrows label septation events.

**Supplemental Figure 5. Increasing agarose concentration reduces out-of-focus hyphal growth.** Brightfield images of hyphae after 17.5 h of growth on either 2% or 5% agarose pads. Asterisks label hyphal tips that grew out of focus. Scale bar, 25 µm.

**Supplemental movie 1. Hyphal growth in *A. pullulans*.** Time lapse Differential Interference Contrast imaging showing hyphal growth (DLY25103). Last frame annotated to show examples of septation (S), branching (Br), and budding from hyphae (B). Time shown in h:min. Scale bar, 50 µm.

**Supplemental movie 2. Polarity in growing hyphae.** Time lapse maximum projection inverted confocal imaging showing localization of polarity sites (mNG-Rac1, black) and nuclei (SpH2B-mCherry, magenta) in growing hyphae (DLY26554) Time shown in h:min. Scale bar, 20 µm.

**Supplemental movie 3. Budding from hyphae.** Time lapse maximum projection inverted confocal imaging showing localization of polarity sites (mNG-Rac1, black) and nuclei (SpH2B-mCherry, magenta) during budding from non-growing hyphal tips and internal compartments (DLY26554). Time shown in h:min. Scale bar, 5 µm.

**Supplemental movie 4. Sec4-GFP accumulation at growing but not non-growing hyphal tips.** Time lapse merged two-color maximum projection confocal imaging showing hyphae with GFP-Sec4 in cyan (vesicle marker) and Psr1$^{1-28}$-mCherry in red (plasma membrane marker) (DLY24564). Time shown in h:min. Scale bar, 20 µm.

**Supplemental movie 5. Tracing hyphal growth with TrackMate.** Time lapse inverted fluorescence image of polarity marker Spa2-mCherry in hyphae (DLY25014). Overlaid TrackMate tracks follow Spa2-mCherry spots as hyphae grow. Tracks and spots colored by track ID. Time shown in h:min. Scale bar, 50 µm.

**Supplemental movie 6. Parasynchronous mitosis.** Time lapse maximum projection confocal imaging of nuclei (SpH2B-mCherry, magenta) and microtubules (3xGFP-Tub2, cyan) in growing hyphae (DLY24568). Arrows point to hyphal compartments that undergo parasynchronous mitosis during the movie. Time shown in min. Scale bar, 20 µm.

**Supplemental movie 7. Pak1 localization in different growth modes.** Time lapse maximum projection inverted confocal images showing localization of polarity sites (Pak1-GFP, black) and nuclei (SpH2B-mCherry, red) in growing cells (DLY25103). (A) A multibudding cell generates multiple polarity sites. This is the same cell shown in Figure 5A. (B) Cells do not polarize during nuclear cycles. These are the same cells shown in Figure 5B. (C) Transient polarization events not associated with polar growth occur before hyphal growth initiates with the formation of a large polarity site. This is the same cell shown in Figure 5C-D. (D) A single large polarity site develops during hyphal initiation. Time shown in h:min. Scale bar, 5 µm.

**Supplemental movie 8. Attempted hyphal initiation in *pak1ΔC* mutants.** Time lapse maximum projection inverted confocal images showing localization of polarity sites (Pak1ΔC-GFP, black) in mutants lacking the Pak1 kinase domain (DLY25650). (A-B) Cells maintain multiple polarity sites. (C-D) Polarity sites can move as travelling waves in these mutants.

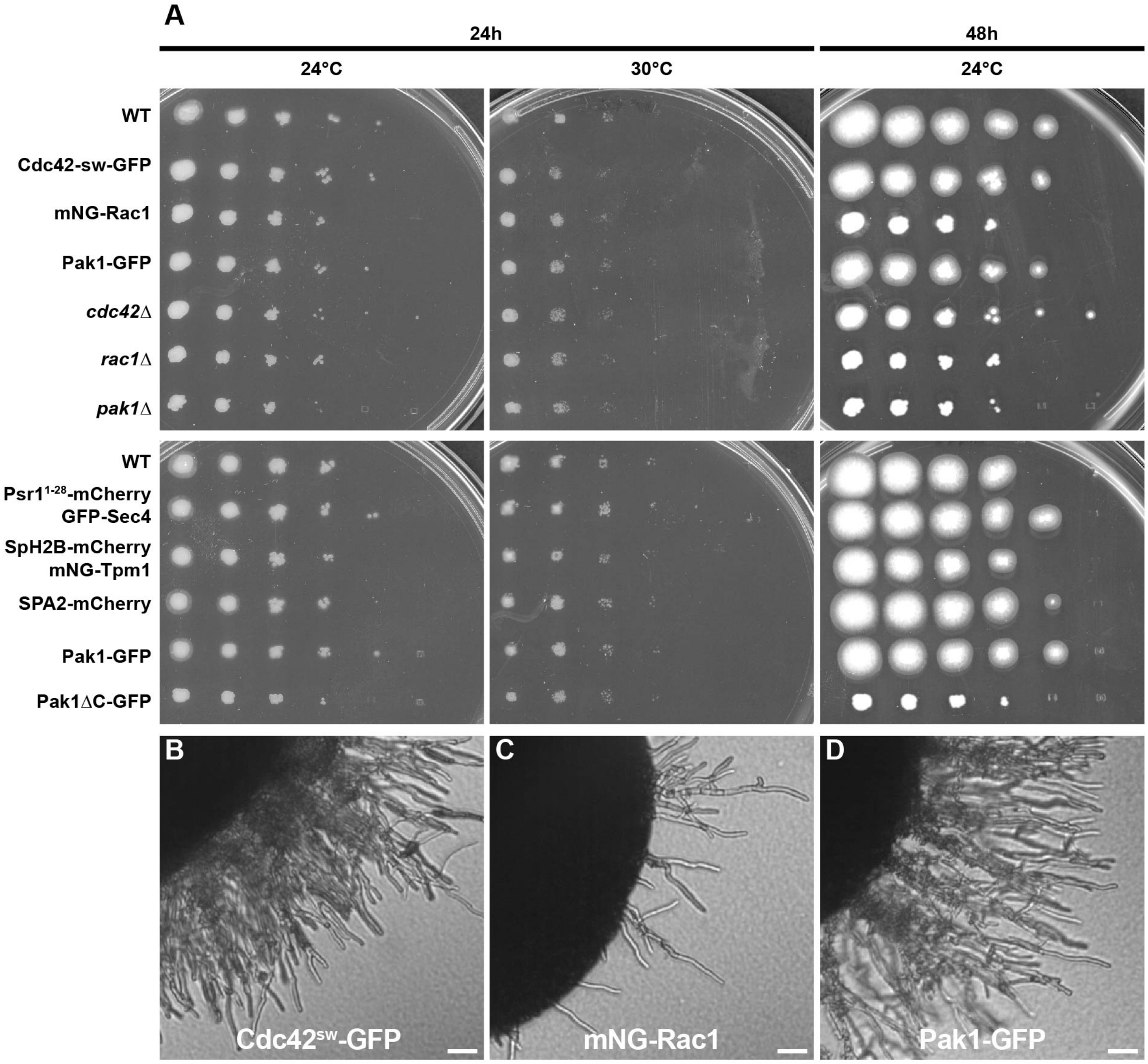


**Supplemental Figure 1. Colony growth and hypha formation of A. pullulans strains used in this study.**

(A) YPD plates with 10-fold serial dilutions of cultures of the indicated strains were grown for the indicated times at the indicated temperatures. Strains from top to bottom in the top plate: DLY23540, DLY26533, DLY26554, DLY25103, DLY25656, DLY25653, and DLY25397 and the bottom plate: DLY23540, DLY24564, DLY26075, DLY25014, DLY25647, and DLY25650. (B-D) Brightfield images of colonies after 2 days of growth: (B) Cdc42sw-GFP (DLY26533), (C) mNG-Rac1 (DLY26554), and (D) Pak1-GFP (DLY25103) Scale bar, 50 μm.

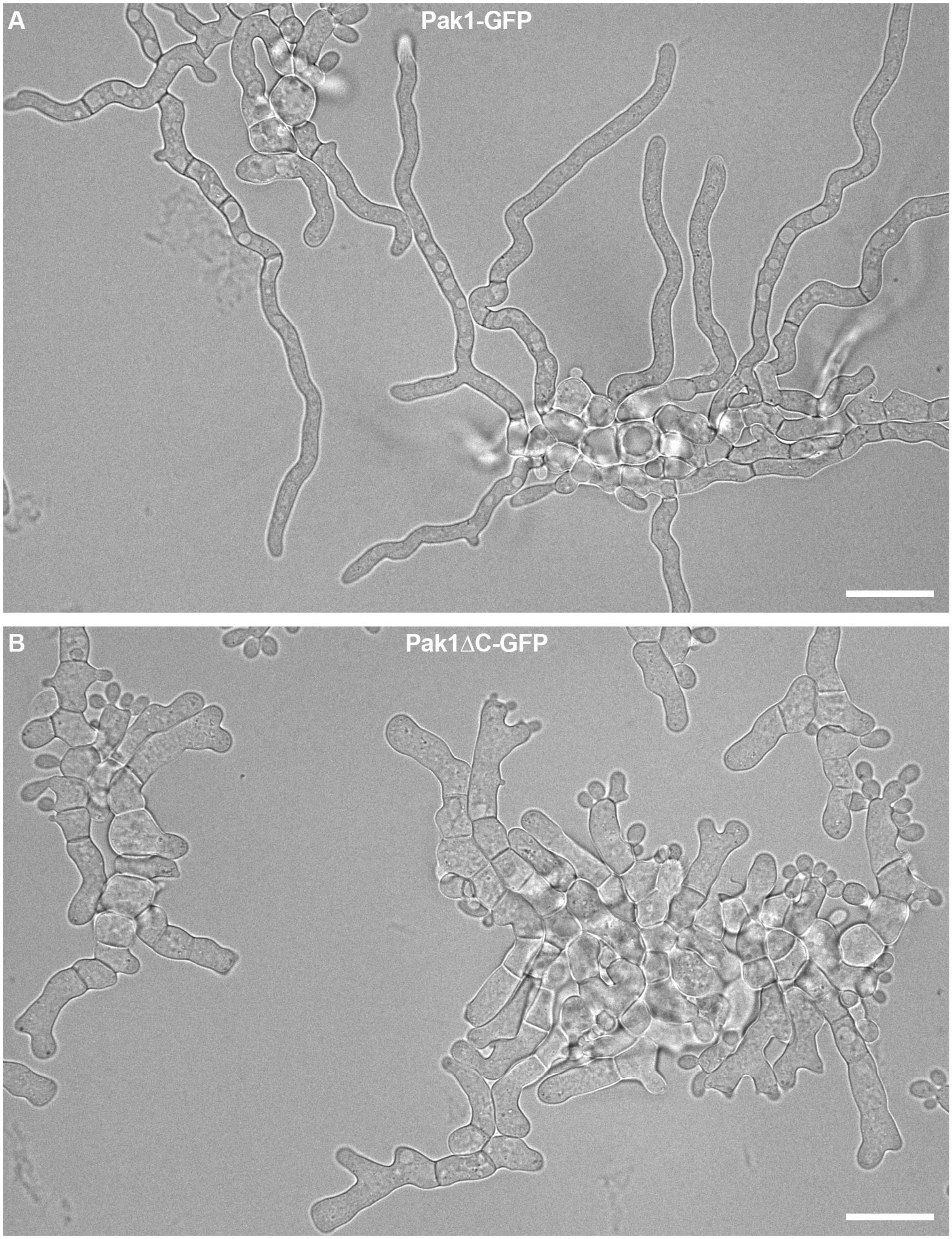


**Supplemental Figure 2. Effect of deleting the Pak1 kinase domain on hyphal development.** Brightfield images of (A) Pak1-GFP (DLY25647) and (B) Pak1ΔC-GFP (DLY25650) mutant cells after 21.5 h of growth under hyphal-inducing conditions. Scale bar, 25 μm.

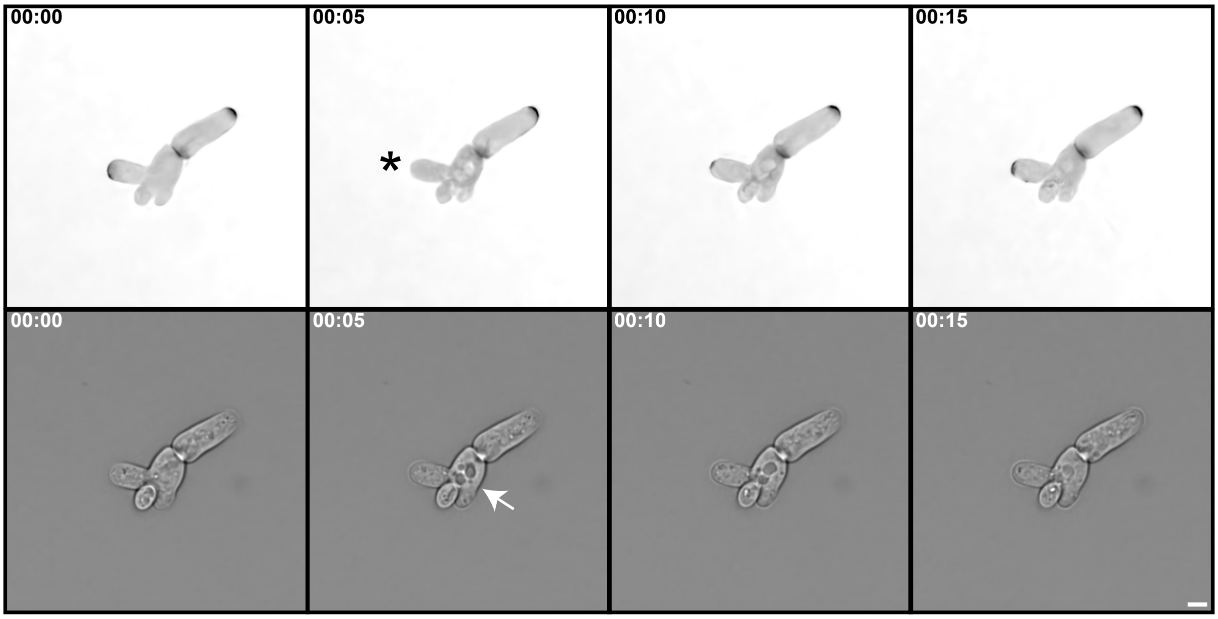


**Supplemental Figure 3. Brief depolarization events in Pak1ΔC mutants.**
Time-lapse inverted fluorescence imaging (top) and Differential Interference Contrast imaging (bottom) of Pak1ΔC-GFP (DLY25650) cells. Asterisk labels transient depolarization and arrow points to vacuoles. Scale bar, 5 µm.

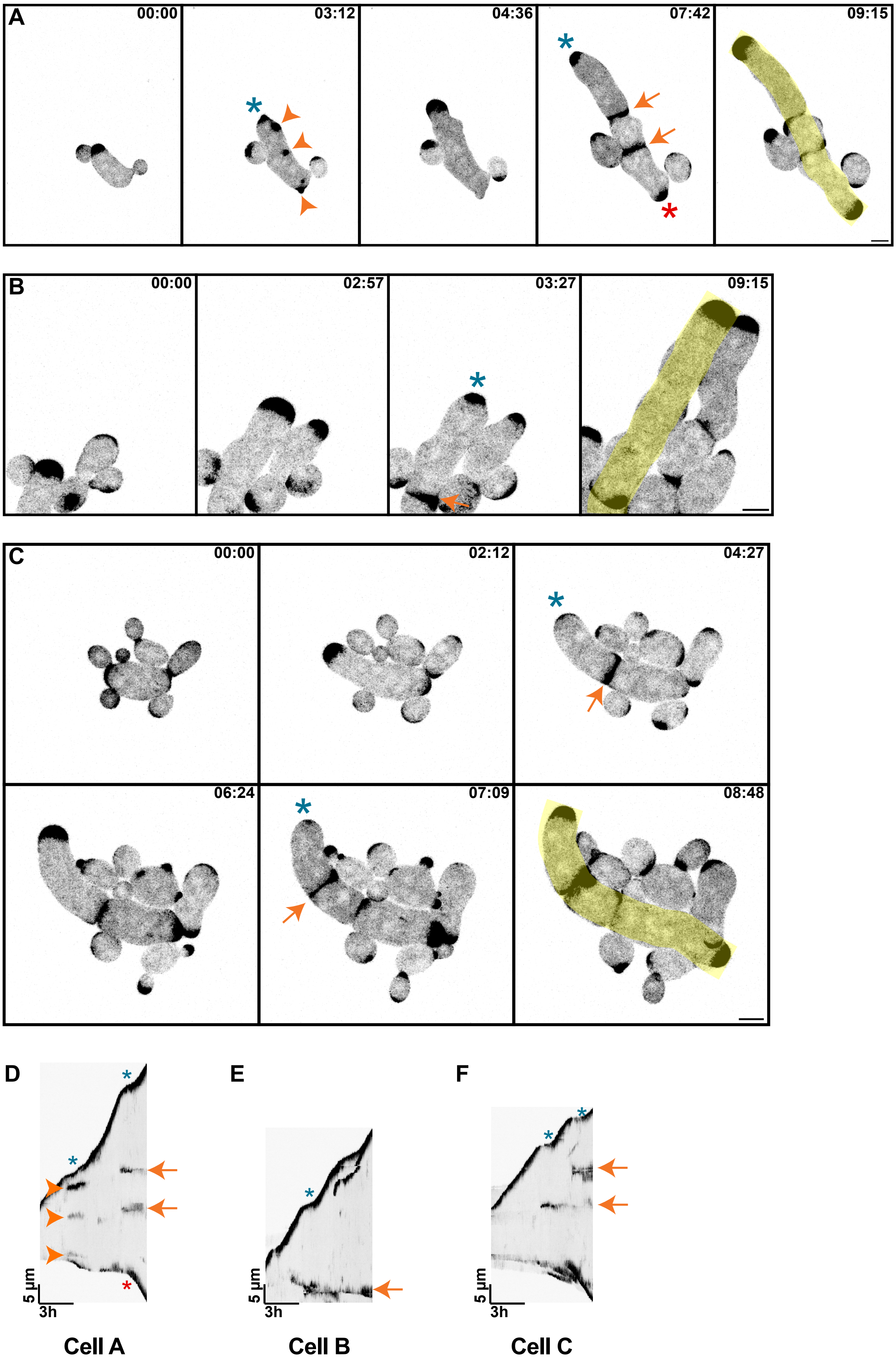


**Supplemental Figure 4. Septation sometimes coincides with growth pausing in Pak1ΔC-GFP mutants.** (A-C) Time lapse maximum projection inverted confocal images showing localization of polarity sites (Pak1ΔC-GFP, black) in mutants lacking the Pak1 kinase domain (DLY25650). Yellow lines were used to create kymographs. Time shown in h:min. Scale bar, 5 µm. (D-F) Kymographs of cells shown in A-C, respectively. Asterisks label growth pauses, arrowheads label additional polarity sites, and arrows label septation events.

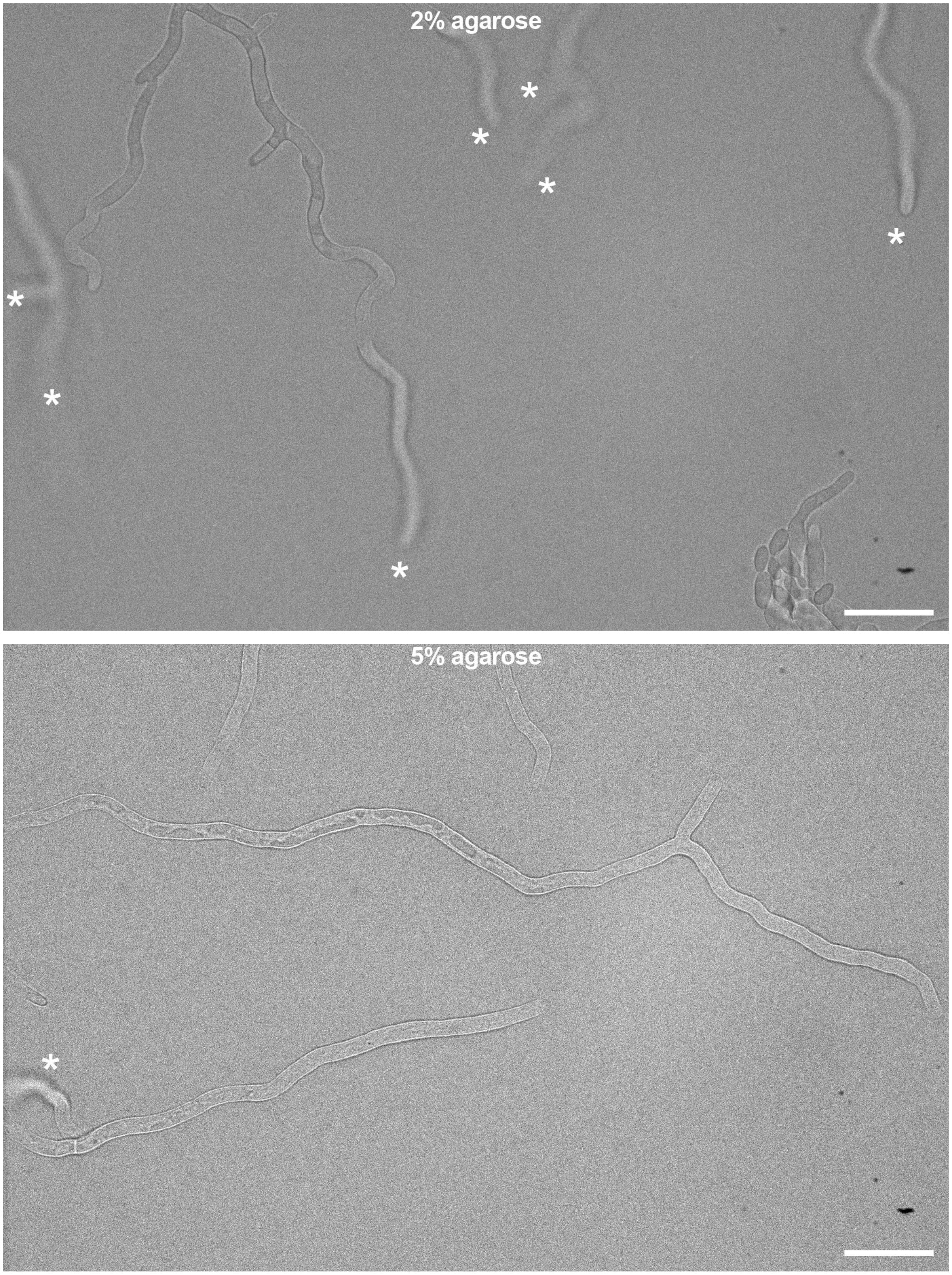


**Supplemental Figure 5. Increasing agarose concentration reduces out-of-focus hyphal growth.** Brightfield images of hyphae after 17.5 h of growth on either 2% or 5% agarose pads. Asterisks label hyphal tips that grew out of focus. Scale bar, 25 µm.
